# Photostimulation of VTA-IC dopaminergic inputs enhances the salience to consolidate aversive taste recognition memory via D1 receptors

**DOI:** 10.1101/2021.11.19.469297

**Authors:** E. Gil-Lievana, G. Ramírez-Mejía, O. Urrego-Morales, J. Luis-Islas, Ranier. Gutierrez, F. Bermúdez-Rattoni

## Abstract

Taste memory involves storing information through plasticity changes in the neural network of taste, including the insular cortex (IC) and ventral tegmental area (VTA), a critical provider of dopamine. Although a VTA-IC dopaminergic pathway has been demonstrated, its role to consolidate taste recognition memory remains poorly understood. We found that photostimulation of dopaminergic neurons in the VTA or VTA-IC dopaminergic terminals of TH-Cre mice increases the salience of a novel taste stimulus regardless of its hedonic value, without altering their taste palatability. Importantly, the inhibition of the D1-like receptor into the IC impairs the salience to facilitate consolidation of an aversive taste recognition memory. Finally, our results showed that VTA photostimulation improves the salience to consolidate a conditioned taste aversion memory through the D1-like receptor into the IC. It is concluded that the dopamine activity from the VTA into IC is required to increase the salience enabling the consolidation of a taste recognition memory. Notably, the D1-like receptor activity into the IC is required to consolidate both innate and learned aversive taste memories but not appetitive taste memory.

## Introduction

Taste memory evolves as a critical system for animal survival through the detection of taste attributes related to the hedonic value, degree of familiarity, and to remember their nutritive or toxic consequences of food to form a memory for future acceptance or avoidance responses (Bermudez-Rattoni, 2004; Scott, 2005). Taste memory involves encoding, storing, and retrieving taste information due to neural plastic changes taking place in a complex network comprising many different brain areas, encompassing the IC, medial prefrontal cortex, basolateral amygdala, nucleus accumbens, and VTA, among others (Yamamoto, 2006, 2008; Ramírez-Lugo et al., 2007; Fontanini et al., 2009; Jezzini et al., 2013).

Several studies have demonstrated that taste learning requires dopaminergic neurotransmission to consolidate the memory representation of tastants (Guzmán-Ramos, 2011; Yiannakas and Rosenblum, 2017). In this regard, the VTA is a vital dopamine supplier that serves a central role in motivating behavior and reward processing. Some evidence suggests that VTA dopaminergic neurons increase their firing rate to signal reward (Berridge and Robinson, 1998; Nomoto et al., 2010; Fiorillo, 2013). However, controversial evidence shows VTA dopaminergic neurons also increase their activity after the presentation of an aversive stimuli (Brischoux et al., 2009; Bromberg-Martin et al., 2010). In this sense, it has been reported that the modulation of the dopaminergic neurons by rewarding or aversive stimuli depends on the brain area(s) to which these dopaminergic neurons project (Lammel et al., 2011; De Jong et al., 2019).

We recently described the functional characterization of the dopaminergic VTA-IC pathway. However, its role in consolidating taste recognition memory remains poorly understood. The photoactivation of TH+ neurons induces electrophysiological responses in VTA neurons, dopamine release, and neuronal modulation in the IC (Gil-Lievana et al., 2020). Importantly, the IC contains the primary gustatory cortex, which serves as a critical structure to consolidate taste recognition memory (Bermudez-Rattoni and McGaugh, 1991; Rosenblum et al., 1993; Chen et al., 2011, Bermudez-Rattoni, 2004; 2014). Accordingly, *in vivo* microdialysis studies show dopamine release triggered in the IC when a novel appetitive or aversive taste is presented (Guzman-Ramos, 2010; Osorio-Gómez et al., 2017). Interestingly, dopamine released during the presentation of novelty enables memory consolidation through the D1-like receptor since post-trial cortical microinjection of D1-like receptor antagonist impedes consolidation of taste recognition memory (Osorio-Gómez et al., 2021). It has been suggested that phasic dopamine activity plays a major role in salience (Bromberg-Martin et al., 2010; Cho et al., 2017) mainly via D1-like receptor (Brenhouse et al., 2008). Salience can be signaled by multiple factors, including intrinsic physical and chemical properties of the stimuli, the association with valenced stimuli, and the physiological state of the organisms, among others (Hyman, 2005; Cowan et al., 2021). Salient stimuli prioritize the consolidation of the relevant over neutral information to drive goal-relevant behaviors (Payne et al., 2008; Moessnang et al., 2012; Alger et al., 2019). We define it as a salience to enable consolidation.

In this work, we aimed to increase the salience of aversive and appetitive taste stimuli through the photostimulation of the VTA-IC dopaminergic pathway to facilitate the consolidation of taste recognition memories. Here we found that the photostimulation of VTA increases the salience to facilitate consolidation of taste stimuli regardless of the natural hedonic value and without altering its taste palatability. Consequently, photostimulation of the VTA dopaminergic terminals into IC also increases the salience to facilitate the consolidation of appetitive and aversive taste stimuli. However, the D1-like receptor activity into the IC is only required to consolidate both innate and learned aversive taste recognition memory.

## Results

### The activity of the VTA dopaminergic neurons increases the intensity of appetitive and aversive taste stimuli

To determine the role of VTA dopaminergic neurons to increase the intensity of innate appetitive and aversive taste stimuli, we tested the performance of mice to consolidate taste recognition memory (TRM) when two different concentrations of saccharin or quinine were presented. We found that a high concentration of saccharin (15 mM, figure 1a) or quinine (504 μM, figure 1b) solution produced a reliable preference for saccharin and avoidance for quinine in mice during the memory test. However, mice did not recognize a very low concentration of either saccharin (5 mM, figure 1a) or quinine solution (126 μM, figure 1b) in comparison to water when presented during the memory test. Therefore, the low concentration of saccharin (5mM) or quinine solution (126 μM) was insufficient to produce a TRM, and they were used as non-salient taste concentrations for our further experiments.

**Figure 1.**
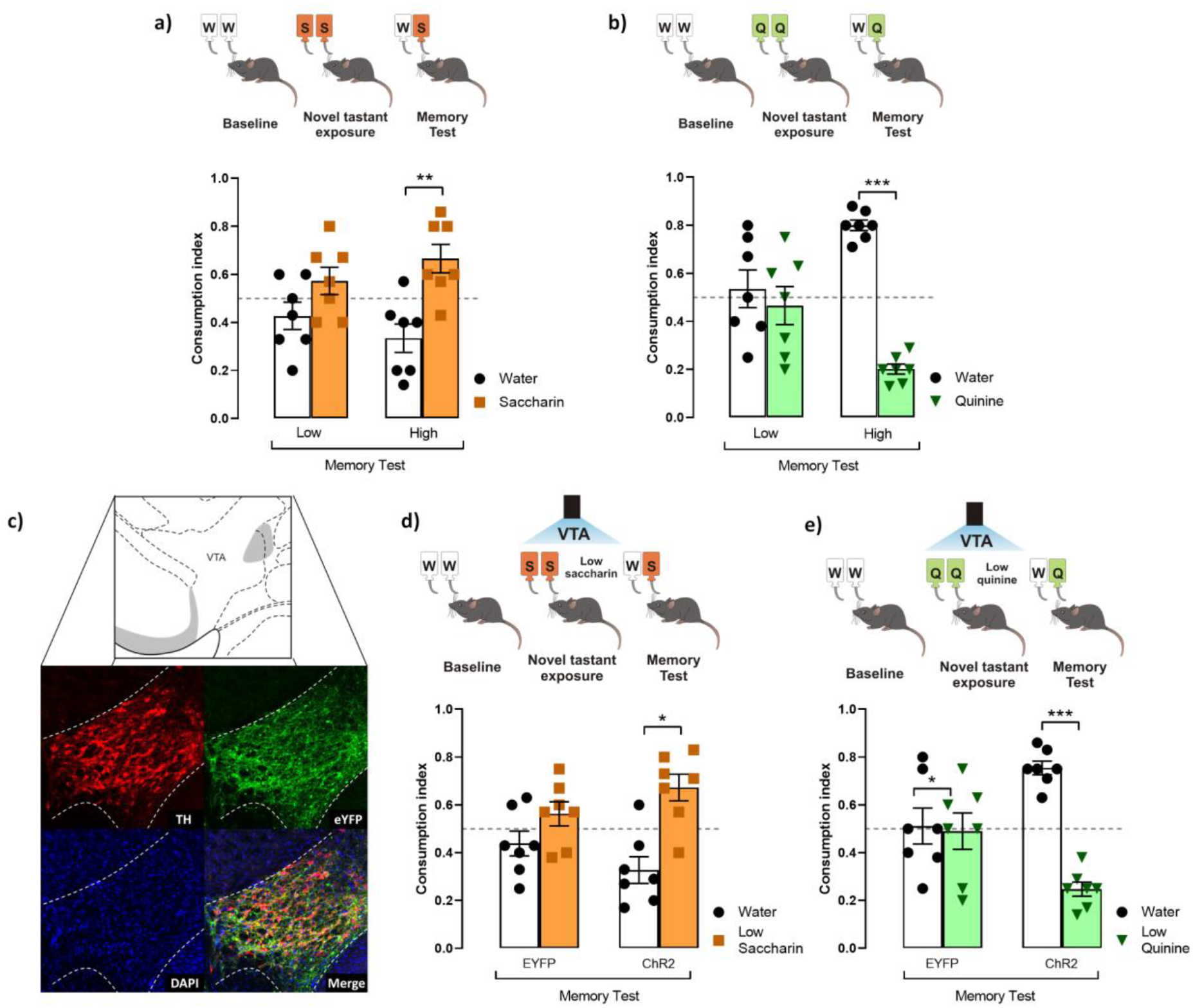
Photoactivation of VTA dopaminergic neurons increases the salience of tastant stimuli regardless of the hedonic value. **a)** Mice were not capable of discriminating low saccharin (5 mM) (n=7) from water during the memory test (paired Student’s *t-test*, t_(6)_=1.283, *p*=0.2470). In comparison, higher saccharin (15 mM) (n=7) was discriminated from water (t_(6)_=2.799, *p*=0.0312). **b)** Accordingly, quinine memory test shows that low quinine (126 μM) (n=7) was not discriminated from water (t_(6)_=0.4440, *p*=0.6726), while higher quinine (504 μM) (n=7) was distinguished from water (t_(6)_=13.78, *p*<0.0001). **c)** Micrographs of the adenoviral infection in VTA. Green shows the reporter protein eYFP, red shows the expression of TH protein, and the last micrograph shows the colocalization between eYFP and TH expression. **d**) Optogenetic stimulation of VTA improves the ability to discriminate saccharin solution from water during the memory test (eYFP group n=7, t_(6)_=1.215, *p*=0.2699; ChR2 group n=7, t_(6)_=3.101, *p*=0.0211). **e)** The photostimulation of VTA improved the performance of mice to discriminate quinine from water during the memory test (eYFP group n=7, t_(6)_=0.1418, *p*=0.8919; ChR2 group n=7, t_(6)_=8.705, *p*=0.0001). \**p*<0.05, \*\**p*<0.01, \*\*\**p*<0.001.

To study whether the VTA dopaminergic neurons increase the salience of appetitive and aversive taste stimuli, we injected TH-Cre mice with an adeno-associated virus encoding Cre-dependent channelrhodopsin-2 protein (ChR2). Enhanced yellow fluorescent protein (eYFP) was used as a reporter protein. Expression of eYFP reporter and tyrosine hydroxylase (TH) immunoreactive neurons in coronal slices of VTA neurons is shown in Figure 1c. Colocalization analysis showed high expression of eYFP in VTA dopaminergic neurons with endogenous tyrosine hydroxylase (TH) immunoreactivity (Figure 1c, merge). We photostimulated the VTA dopaminergic neurons in ChR2 or eYFP mice simultaneously to the presentation of low concentration saccharin or quinine solution during the acquisition session. We found that ChR2 mice showed a reliable TRM, measured by the strong preference for low saccharin solution (figure 1d) or strong avoidance for low quinine solution (figure 1e). However, during the memory test, eYFP mice did not consolidate a TRM to low concentrated saccharin or quinine solutions. These results suggest that VTA dopaminergic neurons increase the salience of low concentration appetitive and aversive taste stimuli to consolidate into TRM.

### Photostimulation of VTA dopaminergic neurons does not alter the palatability of appetitive or aversive tastes

To determine whether the behavioral effects induced by VTA dopaminergic neurons were due to a change in taste palatability, mice were placed in a Brief Access Taste Task (BATT). A BATT measures the oromotor responses (palatability) using the lick rate evoked by tastants during the 5 s reward period (Villavicencio et al., 2018). If dopaminergic neurons are related to taste palatability, we hypothesize that the stimulation would change the lick rate of familiar tastants (García et al., 2021). During this task, mice received either low concentrated saccharin 5mM or quinine 126 μM and water for 5 s per trial (figure 2a). As expected, eYFP mice exhibited a higher lick rate elicited by saccharin and a lower licking rate after quinine delivery (figure 2b). Similar results were also observed when mice were tested at high tastants concentrations (data not shown). Importantly, taste palatability was not altered by the photostimulation of VTA dopaminergic neurons while mice licked for familiar tastes (figure 2c). Specifically, the lick bout duration, a measure of palatability, in the ChR2 mice was not significantly different from eYFP mice (figure 2d; two-way ANOVA, Factor mice, F _(1,21)_=1.188, *p*=0.28). Thus, our data demonstrate that the behavioral effects induced by photostimulation of VTA dopaminergic neurons were not related to taste palatability. Thus, it is more likely that stimulation of DA neurons affected saliency rather than taste palatability.

**Figure 2.**
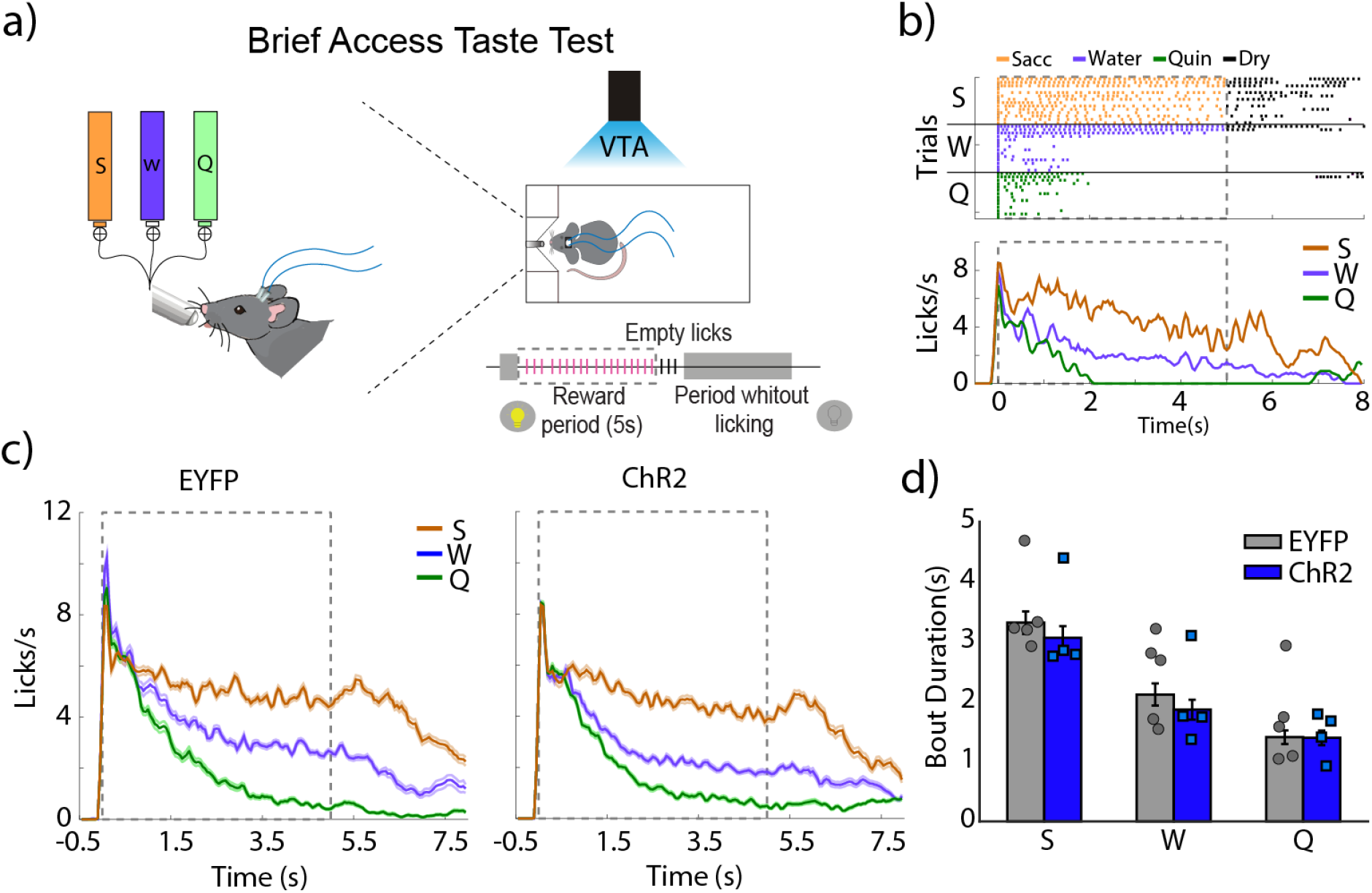
Photoactivation of VTA dopaminergic neurons does not impact taste palatability. **a)** Schematic representation of each trial, in which mice had access to different tastants for a brief time (5 s reward period) to either saccharin, water, or quinine, randomly. Mice received photostimulation during the complete session. **b)** A representative raster plot for licking of one mouse and session. Each tick indicates a single lick and the color the tastant delivered (orange saccharin, blue water, green quinine); black ticks indicate empty (dry) licks. Below is the Peristimulus time histogram (PSTH) from a representative mouse. The dash and gray square indicate the reward period. **c)** Population PSTH for all subjects and sessions. **d)** Mean of lick bout duration (a measure of palatability, i.e., the longer, the more palatable) for each taste during the rewarded period. S, saccharin, W, water, or Q, quinine hydrochloride. No differences in both duration were found between eYFP group (n=5) and ChR2 group (n=4), (two-way ANOVA, Factor mice, F _(1,21)_=1.188, *p*=0.28)

### Specific photostimulation of the VTA-IC dopaminergic terminals increases the salience to consolidate taste stimuli

To determine whether the VTA dopaminergic projections in the IC would solely consolidate taste information (Bermudez-Rattoni, 2014), we photostimulated the VTA dopaminergic projections into the IC (figure 3a) concomitantly with the presentation of novel saccharin (5mM) or quinine (126μM) at low concentrations. ChR2 mice exhibited a reliable TRM during the memory test, as seen by the strong preference for saccharin relative to water (figure 3b) and a strong avoidance for the quinine solution vs. water (figure 3c). Our results showed that the photostimulation of the VTA-IC inputs is sufficient to consolidate a TRM in ChR2 mice but not in eYFP control mice.

**Figure 3.**
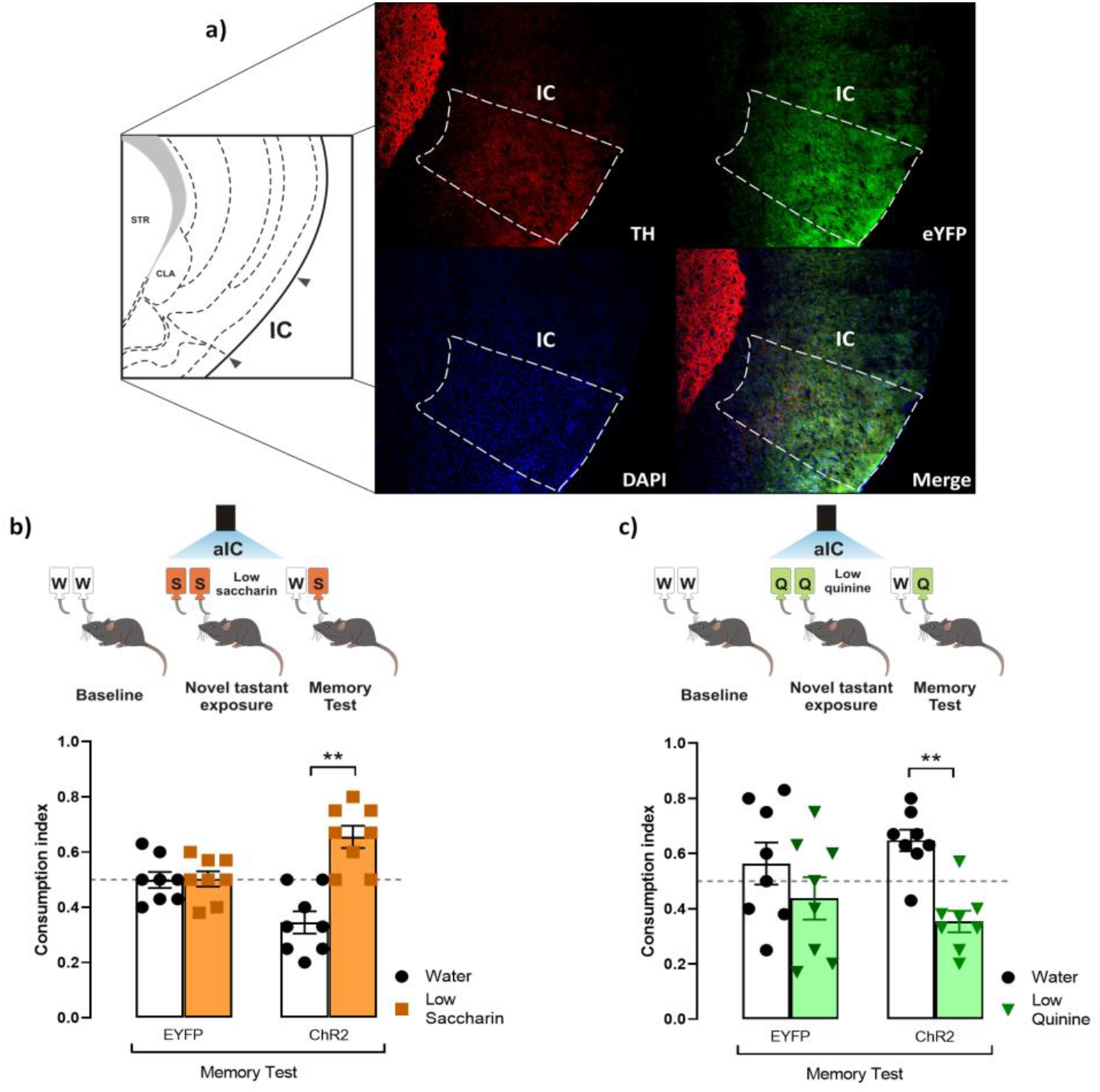
Photostimulation of the VTA-IC dopaminergic pathway increases the salience of taste stimuli. **a)** Representative micrographs of triple immunofluorescence for TH immunoreactive fibers in the IC (top, left), eYFP immunoreactive projections from the VTA into the IC (top, right), DAPI (bottom, left), and merge (bottom, right). **b)** Photostimulation of the VTA-IC dopaminergic projections in ChR2 mice (n=8) enabled the preference for low saccharin solution vs. water (paired Student’s *t-test*, t_(7)_=3.854, *p*=0.0063), but not in eYFP mice (n=8) (t_(7)_=0.06585, *p*=0.9493), during the memory test. **c)** Photostimulation of the VTA dopaminergic projections into the IC led ChR2 (n=8) mice to avoid low quinine solution (t_(7)_=3.767, *p*=0.0070). However, eYFP mice (n=8) did not show a significant difference in consumption between low quinine vs. water (t_(7)_=0.8253, *p*=0.4364). All data are shown as mean ± SEM. \*\**p*<0.01.

### The salience to consolidate aversive but non-appetitive taste recognition memory requires D1-like receptor activity into IC

Having demonstrated the role of dopamine from the VTA into the IC to process appetitive and aversive taste stimuli, we administered an antagonist of D1-like receptors (SCH23390) into the IC before the photostimulation of the VTA dopaminergic neurons in ChR2 and eYFP mice during the presentation of saccharin and quinine solutions. We found that the blockage of the D1-like receptors into the IC did not alter the salience of appetitive TRM (figure 4a), as measured by a preference for saccharin vs. water during the memory test. Nevertheless, the blockage of the D1-like receptors impaired the salience of the aversive TRM (figure 4b). These results suggest that the VTA-IC dopaminergic pathway increases the salience of taste stimuli regardless of the hedonic value to consolidate a TRM. Importantly, the aversive but not appetitive taste stimuli require D1-like receptors into the IC to process the salience of TRM.

**Figure 4.**
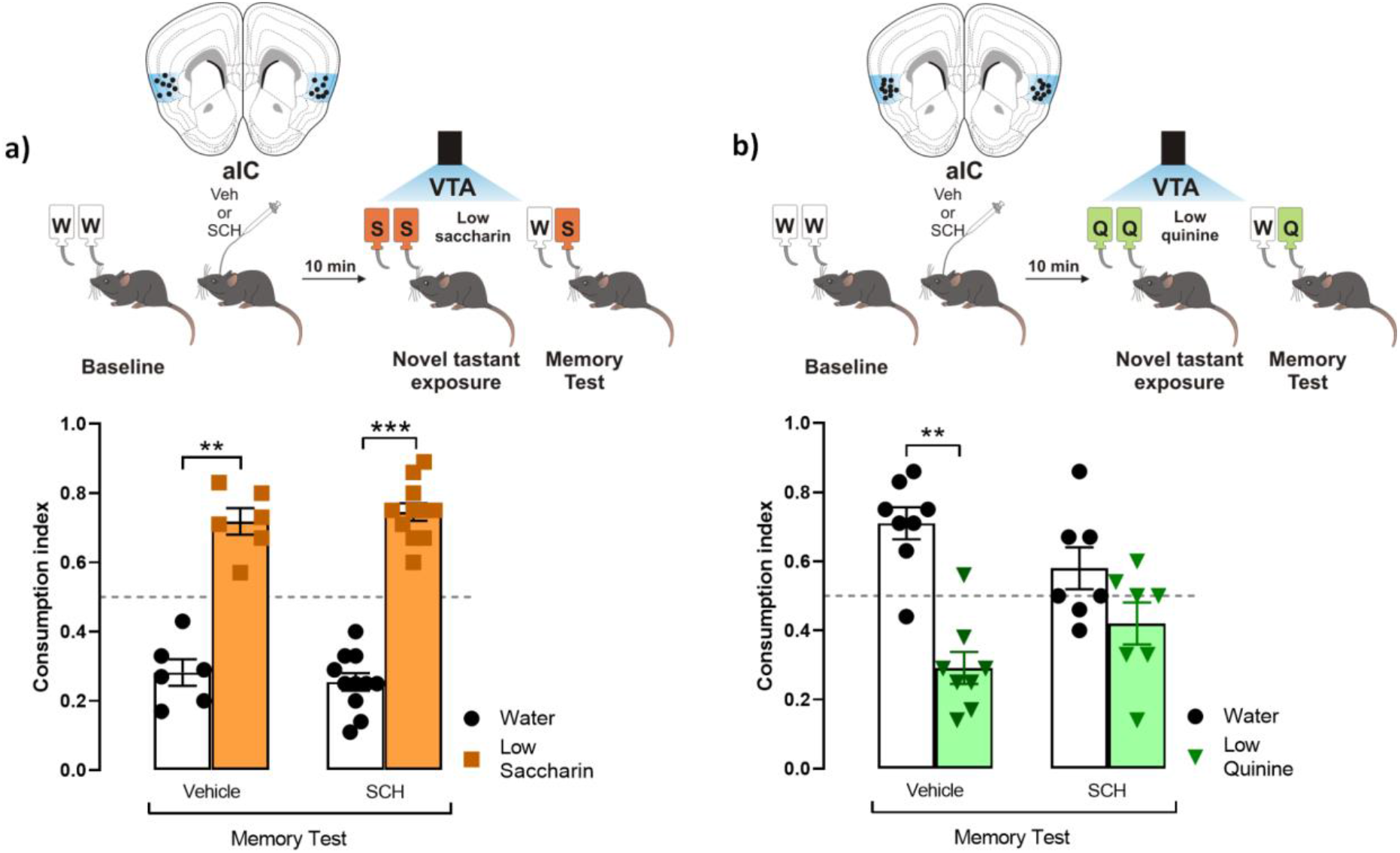
Processing of aversive but not appetitive taste requires D1-like receptor activity into IC. Dopaminergic activity blockade in the IC impedes consolidation of aversive but not appetitive TRM **a)** ChR2 mice were administered with SCH23390 (n=11) or vehicle (n=6) before the photostimulation of the VTA dopaminergic neurons. Both mice groups exhibited a strong preference for the low saccharin solution relative to water (vehicle group paired Student’s t-test, t_(5)_=5.722, *p*=0.0023; SCH group, t_(10)_=9.670, *p*<0.0001). **b)** Administration of SCH23390 before the photostimulation of the VTA-IC dopaminergic pathway impaired the avoidance behavior for the low quinine solution in the ChR2 mice (n=7) (paired Student’s t-test, t_(6)_=1.320, *p*=0.2349) but not in the ChR2 mice administered with vehicle (n=8) (t_(7)_=4.519, *p*=0.0027). All data are shown as mean ± SEM. \*\**p*<0.01, \*\*\**p*<0.001.

### Salience to consolidate conditioned taste aversion requires D1-like receptor activity into IC

Given the VTA-IC dopaminergic pathway’s role in potentiating the salience of an innate aversive taste stimulus to consolidate a TRM through the D1-like receptors. We extended our research to study whether the same pathway is required to process the salience of associative aversive taste memories, such as a conditioned taste aversion memory. First, ChR2 or eYFP mice received the photostimulation of the VTA dopaminergic neurons concomitantly with the presentation of the low saccharin solution, and ten minutes after, they were injected intraperitoneally with a neutral stimulus (NaCl, 66 mg/kg). We found that the eYFP mice did not discriminate between the low saccharin solution vs. water, but ChR2 mice preferred the low saccharin solution from water (figure 5a). Moreover, a low dose of an unconditioned aversive stimulus (LiCl, 48 mg/kg), incapable of inducing a conditioned taste aversion (data not shown), was administered intraperitoneally ten minutes after photostimulation of VTA dopamine neurons and the presentation of saccharin solution. We found that the eYFP mice showed no preference or avoidance behavior for the low saccharin solution vs. water. However, the ChR2 mice showed a robust conditioned taste aversion response due to the aversive association between the gastric malaise produced by the LiCl and the low saccharin solution (figure 5b). Finally, using the protocol described above, ChR2 mice were injected with SCH23390 or vehicle into the IC ten minutes before the conditioned taste aversion. Our results showed that the ChR2 mice injected with the vehicle had a strong conditioned aversive response, with a reliable avoidance behavior for the low saccharin solution associated with the gastric malaise produced by the LiCl. On the other hand, the ChR2 mice administered with SCH23390 were incapable of associating the unconditioned aversive stimulus with the low saccharin solution, showing a preference for the low saccharin solution rather than the water (Figure 5c). In sum, these results suggest that dopamine from VTA into the IC also increases the taste salience enhancing the association of an unconditioned aversive stimulus with an appetitive taste via D1-like receptors.

**Figure 5.**
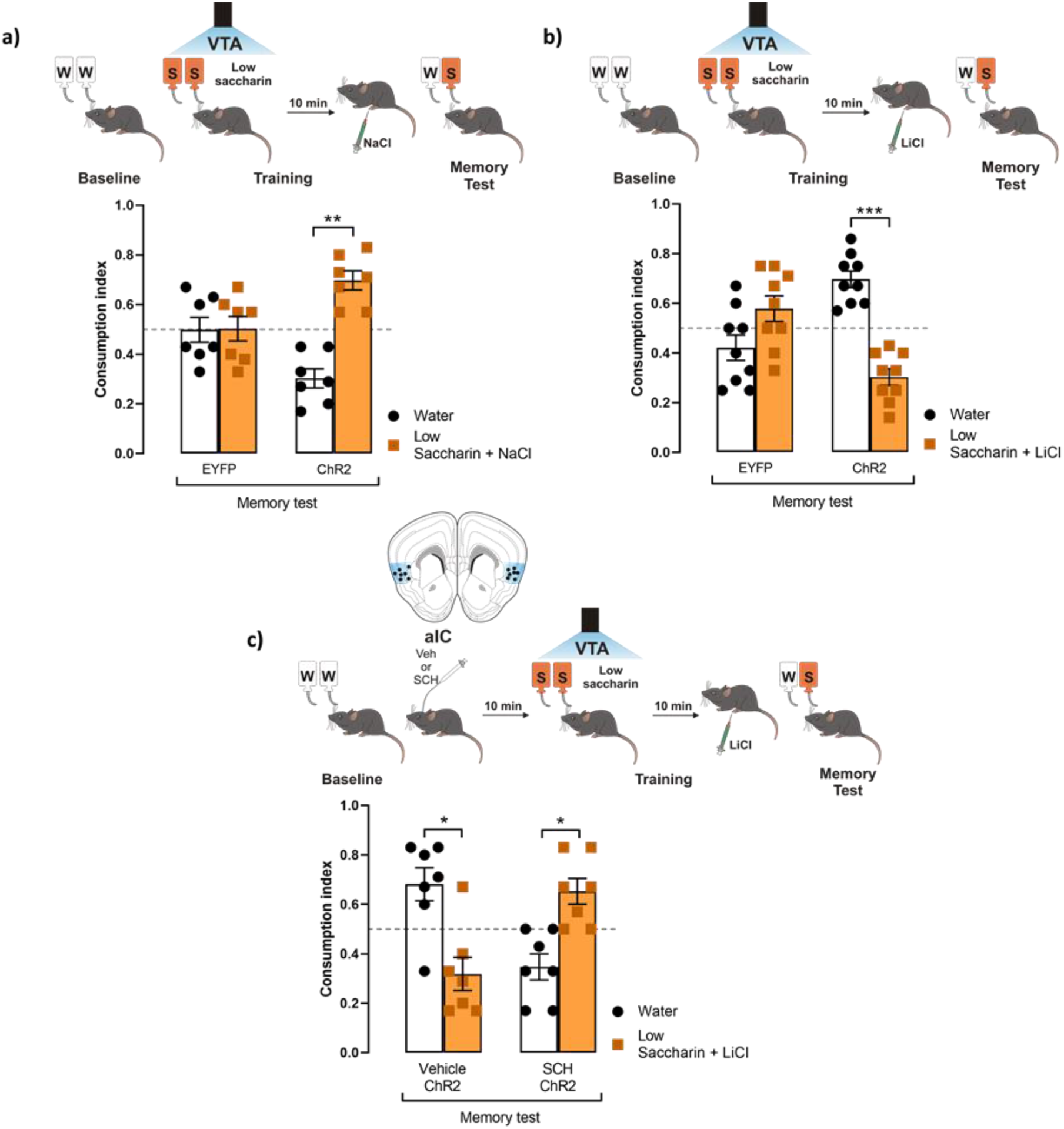
VTA photostimulation increases the salience of conditioned taste aversion for an appetitive taste. **a)** ChR2 (n=7) or eYFP (n=7) mice were photostimulated into VTA when the low saccharin (5 mM) was presented. After, mice were intraperitoneally injected with a neutral unconditioned stimulus NaCl (66 mg/kg). The administration of NaCl in ChR2 mice did not affect the preference for the low saccharin due to the photostimulation of the VTA dopaminergic neurons. **b)** ChR2 (n=9) or eYFP (n=9) mice were photostimulated into VTA when the low saccharin (5 mM) was presented. After, mice were intraperitoneally injected with a low unconditioned aversive stimulus LiCl (48 mg/kg). The administration of LiCl in ChR2 mice decreased the consumption of the low saccharin vs. water, but the consumption of eYFP mice was not affected. **c)** Diagram of coronal slice for IC cannulation in ChR2 mice administered with SCH23390 (n=7) or vehicle (n=7) before the photostimulation of the VTA dopaminergic neurons. After, mice were intraperitoneally injected with a low dose of LiCl. The blockage of the D1-like receptors into the IC impairs the consolidation of the conditioned taste aversion in the SCH group (paired student’s t-test, t_(6)_=2.897, *p*=0.0274). In contrast, the vehicle group showed a reliable conditioned taste aversion (paired student’s t-test, t_(6)_=2.703, P=0.0355). All data are shown as mean ± SEM. \**p*<0.05; \*\*\**p*<0.001.

## Discussion

Our results demonstrate that the VTA increases the salience to facilitate consolidation of a taste recognition memory generated through naturally appetitive or aversive stimuli without an evident alteration on its taste palatability. In this sense, midbrain dopaminergic neurons modulate brain networks through phasic responses to encode not only the novelty, prominence, or surprise of rewarding stimuli but also aversive experiences (Schultz, 1998; Redgrave et al., 1999; Horvitz, 2000; Di Chiara, 2002; Joseph et al., 2003; Pezze and Feldon, 2004; Ungless, 2004; Lisman and Grace, 2005; Redgrave and Gurney, 2006; Schultz, 2007; Moriya et al., 2018). The salience of stimuli, including taste, is adaptive because salient information tends to be preferentially consolidated into long-term memory (Shohamy and Adcock, 2010; Cowan et al., 2021). Importantly, we further extended our previous proposal by demonstrating that the increased VTA dopaminergic neuronal activity did not alter taste palatability. Our results agree with previous results, showing that dopaminergic neurotransmission affects novel taste intake without affecting taste evoked orofacial responses, another parameter of taste palatability (Treit and Berridge, 1990). Therefore, our findings suggest that the dopaminergic activity of the VTA pathway is involved in the formation of the taste memory trace without affecting palatability.

It has been reported that after novel food consumption, there is increased c-fos activity in the VTA and its projection areas within the mesolimbic and mesocortical dopaminergic systems, necessary to form memory (Kest et al., 2012; De la Cruz et al., 2016). Additionally, we recently reported that the photoactivation of the VTA dopaminergic neurons induces dopamine release in the IC, which modulates the activity in this cortical structure (Gil-Lievana et al., 2020). Here we found that VTA-IC dopaminergic activity increases the salience needed to preferentially facilitate consolidation of a taste recognition memory regardless of the hedonic value of taste stimuli. Similar outcomes were obtained by Kutlu and co-workers (Kutlu et al., 2021), who show that dopamine contributes to fit valence-independent perceived salience across different learning paradigms. We hypothesize that the increased salience to facilitate consolidation into IC results from the synaptic potentiation induced by photostimulation of the VTA dopaminergic neurons. It is widely known that tonic and phasic dopaminergic release differentially modifies and modulates the synaptic strength and neuronal activity in many different structures, including the hippocampus, prefrontal, and insular cortices, to serve as a cellular mechanism to establish memory (Matsuda et al., 2006; Sheynikhovic et al., 2013; Rodríguez-Durán and Escobar, 2014; Otani et al., 2015; Navakkode et al., 2017; Papaleonidopoulos et al., 2018). In fact, findings support that brief exposure to a novel environment reduces the threshold to induce long-term potentiation. This facilitatory effect occurs for a short period of time following novelty exposure and depends on the activation of D1-like receptors but is absent in animals that explore a familiar environment (Li et al., 2003). Moreover, it has been demonstrated that aversive stimuli selectively modify synapses on dopaminergic neurons that project to cortical areas (Lammel et al., 2011). Furthermore, the dysfunctional dopaminergic activity alters the synaptic plasticity in the BLA-IC pathway to transform the long-term potentiation into long-term depression associated with memory impairment (Moreno-Castilla et al., 2016).

Although our results show that the VTA-IC dopaminergic activity is sufficient to increase the salience of aversive and appetitive stimuli, we found that D1-like receptor activity is solely necessary to integrate aversive taste information to facilitate the consolidation of an aversive taste recognition memory. Accordingly, the presentation of a novel taste stimulus, regardless of its valence, induces an increment of basal concentration of dopamine in the IC (Osorio-Goméz et al., 2021). However, previous reports have shown that the D1-like receptor is critical for aversive recognition memories formation, increasing neural activity within cortical networks (Heath et al., 2015, Saito et al., 2020). It is parsimonious to hypothesize that the VTA-IC dopaminergic pathway is one of many circuits encoding salience of taste stimuli regardless of the hedonic value due to the processing of salience to consolidate taste information involves multiple brain areas (Bromberg-Martin et al., 2010; Shiner et al., 2015; Cai et al., 2020; Kutlu et al., 2021). However, a D1-like receptor within IC is required for the consolidation and storage of mainly aversive taste recognition memory. Moreover, we found that D1-like receptor activity is also required to establish a conditioned taste aversion. Importantly, the blockage of the D1-like receptor into the IC before a conditioned taste aversion impairs the consolidation, but not the short-term memory, suggesting that the D1-like receptor is involved in the cognitive processing to consolidate the conditioned taste aversion (Osorio-Goméz et al., 2021). This study’s results agree with reports suggesting that D1-like, but not D2/D3 dopaminergic receptor activity is associated with conditioned taste aversion (Fenu et al., 2001).

Dopamine plays a critical role in mediating reward and aversive signals of various stimuli, including visual, auditory, and mechanosensory stimuli (Schultz, 2007; Wise, 2004; Zweifel et al., 2011; McCutcheon et al., 2012; Stelly et al., 2019). We show that although dopamine signaling in the IC is required to consolidate aversive and appetitive taste memories, the downstream molecular events may differ. Indeed, the PKC activity, a kinase involved in neural plasticity processes, is needed in the IC to establish aversive taste memory, but not for appetitive taste memory (Nuñez-Jaramillo et al., 2007). We propose that dopamine from VTA encodes the salience of appetitive and aversive taste, but the interaction of different brain structures (e.g., basolateral amygdala and nucleus accumbens) and other neurotransmitters (i.e., glutamate) within the IC leads to the activation of different signaling pathways to facilitate consolidation of the aversive appetitive nature of the taste memory.

All in all, we found that the activity of the VTA-IC dopaminergic pathway increases the salience to facilitate consolidation of aversive and appetitive taste stimuli. However, the IC only consolidates the aversive taste information through the D1-like dopaminergic receptors. Given the complexity of VTA connectivity, it is reasonable to hypothesize that the encoding of tastant’s salience requires the processing of multiple parallel brain regions. Here we unveiled the VTA-IC dopaminergic pathway as an essential component of this complex circuitry involved in salience to facilitate consolidation of taste recognition memories.

**Table 1.** Total consumption during the memory test. All data are shown as mean ± SEM.

| **Figure 1** | **Volume (mL)** |
| --- | --- |
| a) Low saccharin (n=7) | 1.50 ± 0.11 |
| High saccharin (n=7) | 1.64 ± 0.17 |
| b) Low Quinine (n=7) | 1.47 ± 0.20 |
| High Quinine (n=7) | 1.60 ± 0.13 |
| d) EYFP-Low saccharin (n=7) | 1.64 ± 0.35 |
| ChR2-Low saccharin (n=7) | 1.64 ± 0.21 |
| e) EYFP-Low quinine (n=7) | 1.64 ± 0.17 |
| ChR2-Low quinine (n=7) | 1.57 ± 0.16 |
| **Figure 3** |  |
| b) EYFP-Low saccharin (n=8) | 1.69 ± 0.11 |
| ChR2-Low saccharin (n=8) | 1.44 ± 0.14 |
| c) EYFP-Low quinine (n=8) | 1.47 ± 0.19 |
| ChR2-Low quinine (n=8) | 1.46 ± 0.11 |
| **Figure 4** |  |
| a) Veh-Low saccharin (n=6) | 1.75 ± 0.21 |
| SCH-Low saccharin (n=11) | 1.86 ± 0.11 |
| b) Veh-Low quinine (n=8) | 1.62 ± 0.16 |
| SCH-Low quinine (n=7) | 1.73 ± 0.12 |
| **Figure 5** |  |
| a) EYFP-Low saccharin + NaCl (n=7) | 1.71 ± 0.33 |
| ChR2-Low saccharin + NaCl (n=7) | 1.75 ± 0.11 |
| b) EYFP-Low saccharin + LiCl (n=9) | 1.36 ± 0.12 |
| ChR2-Low saccharin + LiCl (n=9) | 1.47 ± 0.11 |
| c) Veh-EYFP-Low saccharin + LiCl (n=7) | 1.46 ± 0.07 |
| SCH-ChR2-Low saccharin + LiCl (n=7) | 1.57 ± 0.14 |

## Materials and Methods

### Animals

TH-Cre mice (Tyrosine Hydroxylase, FI12 line) express Cre-recombinase protein under the control of the endogenous tyrosine hydroxylase promoter. Breeder mice were kindly donated by Dr. Rui M. Costa from the Champalimaud (Center for the Unknown) and crossed onto C57BL/6J mice for at least six generations. Two-month-old (25-30 g bodyweight) female and male TH-Cre mice were used for all experiments. No difference were found between male and female mice in all experiments. Mice were housed individually at 20 ± 2 °C, 50 ± 5% humidity, under a 12:12 hours light/dark cycle with free access to food and water all time except during behavioral testing. All experiments were conducted during the light phase of the room illumination cycle. All experiments were approved by Instituto de Fisiología Celular (FBR125-18) and complied per the Official Mexican Standard (NOM-062-ZOO-1999).

### Genotyping

The genotyping procedure was previously reported (Gil-Lievana *et al.*, 2020). Once the mice were one month old, a tail snipping procedure was performed, and 1 mm of the tail was removed with sanitized sharp scissors. We used the HotSHOT method for DNA extraction. Briefly, the tail snip was lysed in an alkaline reagent (25 mM NaOH, 0.2 mM disodium EDTA) under heat conditions (95 °C, 1 hour) and further neutralization with a suitable buffer (1 M Tris-HCl, pH 7.4). After centrifugation (2500 rpm, 2 minutes, Hermle Z 233 MK-2), the DNA’s supernatant was recovered. The DNA was used for PCR amplification (201443, QIAGEN). Primers sequences were as follows: Cre forward primer 5’-AGC CTG TTT TGC ACG TTC ACC-3’; Cre reverse primer 5’-GGT TTC CCG CAG AAC CTG AA-3’ (both primers were purchased from Sigma-Aldrich).

### Viral vector

The Cre-inducible adeno-associated virus (AAV) was obtained from the University of North Carolina (UNC) Gene Therapy Center Vector Core. The viral concentrations were as follows: 5.2×10^12^ viral units/ml for rAAV5/EfIα-DIO-hChR2(H134R)-eYFP (ChR2); 6.0×10^12^ viral units/ml for rAAV5/EfIα-DIO-eYFP (eYFP). All viruses were aliquoted and stored at −80 °C until use.

### Stereotaxic surgery

Mice were induced to anesthesia with 3% isofluorane and maintained with 1-1.5% isoflurane (VETone Fluriso^TM^; Matrix VIP 3000, Midmark) until the end of surgery. Once anesthetized, mice were placed in a stereotaxic apparatus (51603, Stoelting) with an incisive adapter (923-B, KOPF instruments). A small incision in the scalp was made, and the head was adjusted to the horizontal plane. The microinjection needles (29-G) were connected to a 10 μl Hamilton syringe and filled with AAV. For all experiments, the mice were bilaterally injected with AAV (0.5 μl) at a rate of 0.1 μl/min with an additional 5 min for diffusion. Mice were implanted with core optic fibers (200 μm) through zirconia ferrules (1.25-mm-wide) in each hemisphere. The AAV was injected into the VTA (from Bregma (mm) AP: −3.08; ML: ±0.60 ML; DV −4.80). The optic fibers were implanted above the VTA (from Bregma (mm) AP: −3.08; ML: ±1.20; DV: −4.30 at 10° angle) or above the IC (from Bregma (mm) AP: +1.40; ML: ±3.30, DV: −3.5). For pharmacological experiments, mice were implanted with bilateral 23-gauge stainless steel cannulas (8 mm long, Small Parts, USA) into IC (from Bregma (mm) AP: +1.40; ML: ±3.30; DV: −3.0). Coordinates were taken from Allens reference atlas of the mouse brain. The cannulas and ferrules were anchored with dental adhesive and dental acrylic cement. Stylets were inserted into guide cannulas to prevent clogging. Mice were allowed to recover for 3 weeks before behavioral procedures.

### Optogenetic stimulation

The conditions for photostimulation were previously reported (Gil-Lievana *et al.*, 2020), briefly: optogenetic stimulation consisted of a diode-pumped-solid-state blue laser (473 nm, 150 mW, 20 Hz; OEM Laser Systems) coupled to 62.5 μm core, 0.22 NA standard multimode hard-cladding optical fiber (ThorLabs) that passed through a single-channel optical rotary joint (Doric Lenses) before being split 50:50 with a fused optical coupler. The intensity of light output was 12-15 mW per split fiber for all experiments.

### Behavioral procedures

Mice were water-deprived only during experimental days. Every afternoon, mice were exposed to bottles of water for 10 min to avoid dehydration. All experiments were conducted during the light phase of the illumination cycle in an acrylic bowl (height 36 cm, diameter 40 cm, CMA 120 bowl, Harvard apparatus). During five consecutive days, two randomized bottles of water were presented for 20 min (baseline). The inclusion criteria were that mice must to consume from both bottles. The mice with a bottle preference were discarded.

#### Taste intensity detection threshold

The next day after baseline, two bottles of quinine (low: 126 μM or high: 507 μM) or saccharin (low: 5 mM or high: 15 mM) were presented for 20 min. Twenty-four hours later, during the memory test, two bottles with water/quinine (low or high) or water/saccharin (low or high) were presented to mice for 20 min.

#### Concomitant optogenetic stimulation and tastant exposure

The next day after baseline, two bottles of 126 μM quinine were presented for 20 min (novel tastant exposure); at the same time, photoactivation of ventral tegmental area (VTA), or VTA projections in the insular cortex (IC) was performed during the 20 min session.

#### Brief access taste task

Mice were water-deprived for 23 h and placed in an operant chamber equipped with a central sipper (Med Associates Inc., VT, USA), where one of three tastants could be delivered (water, quinine 126 μM, or sucrose 5 mM) controlled by a solenoid valve (Parker, Ohio, USA). In each trial, mice randomly received one tastant for 5 s (2 μl drop in each lick), mice decided whether they lick during the entire reward period (García et al., 2021). To start a new trial, mice needed to refrain from licking for a 1-3 s inter-trial interval (ITI) and lick once again after the ITI is finished. Mice were trained during 5 sessions, and 3 additional sessions were performed with laser stimulation, in which mice were opto-stimulated during the entire task at 20 Hz.

#### Conditioned taste aversion and pharmacological manipulations

The next day after baseline, mice were injected into IC with vehicle (0.9 % saline solution), or dopamine D1/D5 antagonist SCH23390 (2 μg/μl, dissolved in 0.9% saline solution, D054, Sigma-Aldrich). Ten minutes after the drug injection, two bottles of 5 mM saccharin were presented for 20 min; simultaneously, photoactivation of VTA was performed during the 20 min session (the photostimulation conditions were like those previously described). Ten minutes after the tastant exposure, mice received an intraperitoneal injection of 0.15 M LiCl at 48 mg/kg body weight or 0.15 M NaCl at a dose of 66 mg/kg body weight.

Twenty-four hours after the novel tastant exposure, a memory test was performed in all cases; one bottle that contained the taste (used during the novel tastant exposure/training) and one bottle of water were presented to mice for 20 min. Consumption indexes for the test were calculated by dividing the volume of the tastant by the total volume consumed: tastant / (tastant + water).

### Immunofluorescence

After the test, mice were sacrificed with an overdose of intraperitoneal pentobarbital monosodium (200 mg/kg). Intracardiac perfusion was performed with 0.9% saline solution and pre-fixed with 4% paraformaldehyde in 0.1 M phosphate buffer solution. Brains were removed and fixed in 4% paraformaldehyde and stored for one week. Brains were treated with 30% sucrose at least 2 days before slicing. Brains were sliced in 40 μm sections using a cryostat (Leica, CM1520). Free-floating sections were incubated with anti-TH (1:1000, rabbit, P40101, Pel-Freez, Rogers, AR) overnight, at 4 °C. Sections were washed with trizma buffer solution added with a triton (TBST; 150 mM NaCl, 100 mM trizma base, 0.1% triton X-100; all purchased from Sigma-Aldrich) incubated with CY3-conjugated goat anti-rabbit (1:250, AP132C, Millipore, Darm-Stadt, Germany) for 2 hours. Antibodies were incubated in 5% bovine serum albumin in TBST. Sections were washed with TBST and incubated with 300 nM 4‘,6-diamidino-2-fenilindol (DAPI, D9542, Sigma-Aldrich) for 1 minute. After a final wash with TBST, sections were mounted in Dako fluorescence mounting medium. Immunofluorescence was observed using a *ZEISS LSM 800* confocal microscope.

### Statistical analysis

For experiments, we used the minimum sample necessary to obtain a mean and standard deviation required for a Cohen’s d parameter equals or greater than 0.8 (Lakens, 2013). Data were analyzed using GraphPad Prism software version 8.0 and Matlab R2021a. The Kolmogorov-Smirnoff test was performed for normal distribution. Data were plotted as mean ± SEM. Paired Student’s *t-*test and two-way ANOVA were used. For all statistical analyses *p*<0.05 threshold was considered statistically significant.

## Acknowledgments

We thank Dr. Rui M. Costa and Dr. Fatuel Tecuapetla for the TH-Cre mice. We thank Dr. Daniel Osorio Gómez for their comments on an earlier version of the manuscript. Dr. Luis Rodríguez Durán, Dr. Josué O. Ramírez Jarquín and Cecilia Acevedo Huerta for technical assistance. Dra. Ruth Rincon Heredia and Dr. Abraham Rosas Arellano for imaging support. This work was supported by Productos Medix 3247, Cátedra Marcos Moshinsky (to R.G.), and the CONACyT grants 250870, FOINS 474 and DGAPA-PAPIIT-UNAM IN212919 to F.B-R.

